# Updating memories of unwanted emotions during human sleep

**DOI:** 10.1101/2022.07.18.500414

**Authors:** Tao Xia, Ziqing Yao, Xue Guo, Jing Liu, Danni Chen, Qiang Liu, Ken A. Paller, Xiaoqing Hu

## Abstract

Post-learning sleep contributes to memory consolidation. Yet, it remains contentious whether sleep affords opportunities to modify or update emotional memories, such as those people would prefer to forget. Here we attempted to update memories during sleep using spoken positive emotional words paired with cues to recent memories for aversive events. Affect updating using positive words during human non-rapid-eye-movement (NREM) sleep, compared with using neutral words instead, reduced negative affect judgments in post-sleep tests, suggesting that the recalled events were perceived as less aversive. EEG analyses showed that emotional words modulated theta and spindle/sigma activity. Specifically, to the extent that theta power was larger for the positive word than for the following memory cue, participants judged the memory cues less negatively. Moreover, to the extent that sigma power was larger for the emotional word than for the following memory cue, participants showed higher forgetting of unwanted memories. Notably, when the onset of individual positive word coincided with the upstate of slow oscillations, a state characterized by increased cortical excitability during NREM sleep, affective updating was more successful. In sum, the affect content of memories was altered via strategic spoken words presentations during sleep, in association with theta power increases and slow-oscillation upstates. These findings offer novel possibilities for modifying unwanted memories during sleep, without requiring conscious confrontations with aversive memories that people would prefer to avoid.

## Introduction

Sleep sculpts our emotional memories via offline consolidation (Goldstein & Walker, 2014; Rasch & Born, 2013; Talamini & Juan, 2020; Walker & van der Helm, 2009). But whether memories can be updated and modified during sleep? Unwanted memories, such as for traumatic or shameful experiences, can be particularly debilitating for cognitive functioning and mental well-being (Dunsmoor et al., 2022; Pitman et al., 2012). However, controlling unwanted memories can be daunting, given the challenge of top-down cognitive control abilities (Anderson & Hulbert 2021; Hu et al., 2017). Moreover, people often wish to avoid thinking of such unwanted memories, thereby precluding direct confrontation and control. It would therefore be desirable to modify unwanted memories without direct confrontation and without engaging cognitive effort. Here, we examined the novel hypothesis that unwanted memories can be updated during sleep, bypassing the challenge of confronting a negative memory.

An established paradigm to manipulate memory processing during sleep is known as targeted memory reactivation (TMR, Paller et al., 2021; Rasch et al., 2007; Rudoy et al., 2009). Auditory or olfactory stimuli are first linked with awake learning, and then specific memories are reactivated during sleep via unobtrusive presentations of those stimuli, which are especially effective during non-rapid eye movement sleep (NREM sleep, for a meta-analysis see Hu et al., 2020). TMR has been shown to influence many types of memory, including spatial memory, motor memory, emotional memory, linguistic memory, and others (Ai et al., 2015; Antony et al., 2012; Cariney et al., 2014; Cheng et al., 2021; Schechtman et al., 2021; Schreiner et al., 2015; Wassing et al., 2019; see Paller et al., 2021 for a review). Notably, researchers also adapted TMR to modify fearful or emotional memories during sleep (Ai et al., 2015; Ashton et al., 2018; Cairney et al., 2014; Hauner et al., 2013; He et al., 2015; Hutchison et al., 2021; Lehmann et al., 2016; Pereira et al., 2022; van der Heijden et al., 2022), but the results to date are mixed; TMR either weakened, strengthened, or had null effects on emotional memories. This evidence does not provide convincing support for the idea that these TMR methods can effectively update unwanted memories.

However, instead of provoking reactivation by presenting a stimulus linked to emotional information from the pre-sleep learning phase, some investigators have used more complex sleep learning or TMR paradigms. These paradigms involved an attempt to form new associations during sleep, such as odor-odor, odor-tone or even word-word associations (Arzi et al., 2012; Koroma et al., 2022; Züst et al., 2019). In one study, such procedures even reduced the unhealthy habit of cigarette smoking (Arzi et al., 2014). In a particularly relevant TMR study, Simon and colleagues (2018) trained tones to be associated with efforts to forget, and then played those forgetting-associated tones following memory cues during sleep. Results showed forgetting of episodic memories. Episodic forgetting was also shown with a related procedure in which forgetting-associated tones were played during sleep after directed forgetting was attempted prior to sleep (Schechtman et al., 2020).

Another variation on the TMR paradigm, yet to be explored, is to attempt to update a memory by combining a memory cue with a stimulus of opposite valence (i.e., counterconditioning). During wakefulness, research in fear learning or evaluative conditioning has shown that counterconditioning can be an effective procedure to change emotional responses and even maladaptive behaviors (Hu et al., 2017; Keller et al., 2020; Van Gucht et al., 2010). These counterconditioning research, combined with sleep learning and TMR, raise a novel possibility that via pairing positive emotional stimuli with memory cues during sleep, people may update the affect tone of previously learnt aversive memories.

Here, we designed a novel, sleep-based memory updating procedure to test the extent to which we can update unwanted memories via pairing positive words with memory cues. The task consists of three sessions: pre-sleep learning, sleep-based updating, and post-sleep testing. Prior to sleep, participants learned cue-target pairings involving initially neutral pseudoword cues and aversive emotional pictures. We also included memory tests following the learning procedure. During post-learning NREM sleep, we unobtrusively played spoken emotional words (positive or neutral) as unconditioned stimuli, followed by the memory cues. We presented spoken emotional words immediately before the memory cues to ensure that the effects are due to affective conditioning, instead of memory disruption that might result if words were played following memory cues. To assess the behavioral effects of sleep-based updating, we measured participants’ affective judgments and accuracy of unwanted memories after sleep. We hypothesized that by repeatedly pairing positive emotional words with memory cues during NREM sleep, participants’ negative affective responses toward cues would be weakened in the post-sleep tests.

We further hypothesized that if unwanted memories can be updated, stimulus-elicited brain activity during NREM sleep would be critical for updating to emerge and may be observable in EEG measures. TMR and sleep-learning studies pinpoint the role of theta (5-9 Hz EEG signals) in emotional prosody processing, emotional memory reactivation/reinstatement, and encoding of emotional stimuli during NREM sleep (Arzi et al., 2012, 2014; Blume et al., 2017; Canales-Johnson et al., 2019; Legendre et al., 2022; Lehmann et al., 2016). We thus hypothesized that the valence of the spoken word could modulate theta power during NREM sleep, which will then drive successful affective updating.

In addition to theta power, spindle activity within sigma band (12-16 Hz) and slow-wave activity within delta band (0.5-4 Hz) are instrumental for sleep-mediated memory reactivation and consolidation. Specifically, spindle and spindle-related sigma power have been associated with information processing and emotional memory reactivation during NREM sleep (Andrillon et al., 2016; Andrillon & Kouider, 2020; Legendre et al., 2022; Lehmann et al., 2016). Here, given that we presented pairs of stimuli during sleep, we were interested in whether sigma power elicited by the emotional words and the memory cues would influence post-sleep memory. In addition, the cortical slow oscillation (SO, 0.5-2 Hz), a defining neural oscillation of deep sleep, encompasses downstates and upstates that reflect neural hyperpolarization and depolarization, respectively. The SO upstate is thought to comprise a transient period suitable for high-level cognitive processing and long-distance cross-region communication (Destexhe et al., 2007; Niknazar et al., 2022; Schabus et al., 2012). Indeed, when cues or auditory stimulation were played during SO upstates in particular, stronger memory benefits or sleep learning effects emerged (Göldi et al., 2019; Ngo et al., 2013; Züst et al., 2019). Here, we focused on the contingency between SO phase and onset of spoken words and examined whether such contingencies influence post-sleep affective updating.

## Methods

### Participants

Forty-six native Chinese speakers participated in the study. Participants reported regular sleep-wake cycles, did not take any medication that impair sleep or mood, had no history or current diagnosis of neurological or psychiatric illnesses. Participants were compensated with monetary incentive. Six participants were excluded because they reported hearing the words while sleeping, four participants were excluded because they had fewer than 48 pairing trials (i.e., one pairing block), and five participants dropped from the experiment before sleep. One participant’s sleep EEG data were not saved due to equipment breakdown. The final analyses included 31 valid participants in the behavioral analysis (Gender: 9 male, 22 female, Age: Mean ± SD., 21 ± 2) and 30 valid participants in the EEG analysis (with at least 48 pairing trials, Mean ± S.E., Positive: 191 ± 10; Neutral: 188 ± 10; *t*(29) = 1.70, *p* = .100). The study was approved by the Human Research Ethics Committee of the University of Hong. All participants provided written consent prior to participation.

### Materials

Thirty-six two-syllabi pseudowords were created by randomly pairing two neutral characters. We then selected 12 positive words (Valence: Mean ± S.D., 7.00 ± 1.28; rating obtained from 9-point Likert-scale, with 1 extremely negative to 9 extremely positive) and 12 neutral words (Valence: Mean ± S.D., 5.29 ± 1.58) from the Chinese Affective Word System (Wang et al., 2008). Vocalization of the pseudowords, positive and neutral words were generated via Text-To-Speech of iFLYTEK (word duration, Mean ± S.D., 761 ± 101 ms). For emotional pictures, we selected 36 negative pictures (Valence: Mean ± S.D., 3.14 ± 0.53; Arousal: Mean ± S.D., 4.43 ± 1.22; ratings obtained from 9-point Likert-scale, with 1 extremely calm down to 9 extremely excited). These pictures are from three categories: Animal, Baby, Scenes, with each category containing 12 pictures (sources: International Affective Picture System, IAPS, Lang et al., 1997, Nencki Affective Picture System, NAPS, Marchewka et al., 2014, and from Internet)

### Procedure

Participants completed the following three sessions: 1) pre-sleep learning in which they learned cue-target pairings to acquire negative emotional memories; 2) sleep-based affective updating in which they were played with spoken word-cue pairings during NREM sleep; 3) post-sleep tests in which they were tested on affect responses and memory promoted by cues.

In the pre-sleep learning session (∼20:30), participants completed the following tasks in order: 1) affect rating of negative pictures; 2) cue-target learning involving pseudoword as cues and negative pictures as targets; 3) baseline affect judgment task; 4) baseline cue affect rating; 5) baseline cued recall. Participants went to sleep (∼23:00) after completing these tasks.

Participants first rated each of the 36 negative emotional pictures on valence (1 extremely negative to 9 extremely positive Likert scale) and arousal (1 extremely calm to 9 extremely excited Likert scale). Each trial started with an 800-ms fixation, followed by pictures being presented on the center of the screen until participants gave responses using a computer mouse. Pictures from all three categories (Animal, Baby, Scene) were randomly presented.

During cue-target learning, participants memorized 36 pseudowords-negative picture pairings via four viewing and test-feedback rounds. In the viewing phase, each trial started with a fixation (jittered 800-1200ms), followed by two-syllabus aurally presented pseudowords (∼1000ms). After a 1000-ms blank screen, a pseudoword-picture pairing was presented for 1500 ms on the center of the screen, while the spoken pseudoword was played again. After participants viewed all 36 pseudowords-picture pairings, they took a 1-minute break, followed by a test-feedback phase. Here, participants were visually and aurally presented with the pseudoword (∼1000 ms), together with three pictures being presented on the screen. Participants were prompted to identify the correct picture that was paired with the spoken pseudoword from the previous viewing session. Note that all pictures in this test-feedback phase were chosen from the viewing session, preventing participants from relying on familiarity to make a correct judgment. Upon participants’ choice, a “correct” or “incorrect” feedback was provided regardless of accuracy, followed by the presentation of correct pseudoword-picture pairing for 1500 ms. Participants were presented with their recognition accuracies at the end of each test-feedback phase. Participants underwent this viewing and recognition-feedback round for four times.

In the affect judgment task, each trial started with an 800-1200ms fixation, followed by the cues being played aurally and visually on the center of the screen for ∼1000 ms. Participants made a speedy negative or neutral affect judgment towards the cue, using the left vs. right keys within 1.5s. Participants completed this task before and after sleep to measure the affective updating effect.

In the cued recall tasks, each cue started with a fixation (800-1200ms), followed by the cue word being aurally presented for ∼1000ms. Participants were asked to verbally describe the paired pictures in as much detail as possible within 15s. The inter-trial-interval was set to be 3s. Participants completed this cued recall task before and after sleep to measure memory changes.

Participants woke up around 7:00 the next morning, and completed the following task in order: 1) affective judgment task; 2) affect rating; 3) cued recall.

### Sleep-based affect updating

We randomly selected two out of three categories of negative memories and their associated memory cues to be paired with positive (one category, 12 cues) or neutral (one category, 12 cues) words during sleep. The remaining one category (12 cues) were assigned to the non-pairing condition, i.e., they were not paired with any words during sleep. One of the categories was randomly selected and paired with positive words, while another category was paired with negative words. Memory categories assigned to positive pairing, neutral pairing and non-pairing conditions were counterbalanced across participants.

Participants went to bed around 23:00. Well-trained experimenters started playing the spoken words when participants entered slow-wave sleep for at least 5 minutes. Each pairing trial started with a spoken positive or neutral words (∼1s), followed by a spoken pseudoword i.e., memory cue (∼1s) with an ISI of 1 s. The ITI was 4s. Each pairing block contained 24 trials that were randomly presented, with 12 positive words + cue trials and 12 neutral words + cue trials. Participants took a 1-minute break between blocks. After every three pairing blocks, a non-pairing block was played. In the non-pairing block, all 36 cues (12 positive pairing cues, 12 neutral pairing cues, 12 non-pairing cues) would be played randomly without any paired words. The ITI was 4s. Each round included three pairing blocks and one non-pairing block. Playing was paused if participants entered REM or N1 sleep or show arousal or wake (e.g., burst of EMGs, alpha activity). The experimenter would end the procedure when 1) seven rounds (i.e., 21 pairing blocks and seven non-pairing block) were completed or 2) at 2:00 in the morning, whichever came first.

### Equipment

All experimental tasks were implemented with Psychopy 3.0 (Peirce, 2007). During sleep, all aurally presented stimuli were played via a loudspeaker (∼47-dB sound pressure level) mounted one meter above the bed, with white noise being played throughout the night.

### EEG recording and preprocessing

Sleep EEGs were recorded using a 64-channel EEG cap connected to an eego amplifier (ANT neuro), with electrodes mounted according to the International 10-20 system. F3/F4, C3/C4, P3/P4, and O1/O2 were selected for online sleep monitoring. One EOG channel was placed below the left eye to monitor eye movements. Two additional bipolar EMG electrodes were placed on the chin to record EMG. On-line EEG data were bandpass filtered from 0.5 to 40 Hz at a 500 Hz sampling rate.

We used MNE-Python for offline EEG pre-processing (Gramfort et al., 2013). First, EEG data were down-sampled to 200 Hz. Second, EEG data were filtered with a bandpass of 0.5-40 Hz. Third, bad channels were visually identified and marked. Next, data were re-referenced to the average of all non-marked electrodes after removing the M1 and M2. Fifth, for trials in the pairing blocks, continuous EEG data were segmented into short (-1.5s to 5.5s) and long (-15s to 15s) epochs relative to the onset of the spoken word. We used the short [-1.5 - 5.5s] 7s epochs in stimulus-locked event-related potentials (ERPs) and time-frequency analyses, and the long [-15 - 15s] 30s epochs in stimulus-locked sleep event detection analyses on a trial basis. For trials in the non-pairing blocks, continuous EEG data were segmented into [-1.5s – 3.5s] 5s epochs relative to the onset of memory cues. Lastly, artifacts were visually inspected and deleted, followed by bad channel interpolation.

### Behavioral analysis

For behavioral data, we focused on affect changes from pre- to post-sleep affect responses. Specifically, for the affect judgment task, we calculated affect judgment changes by subtracting the pre-sleep baseline neutral response ratio from the post-sleep neutral response ratio. A higher change score, i.e., more neutral judgments or fewer negative judgment from pre- to post-sleep, would indicate higher affect changes toward neutrality. For the affect rating task, we similarly calculated affect rating changes by subtracting pre-sleep baseline valence/arousal ratings from post-sleep ratings. A higher valence/arousal change score would indicate more positive/arousal changes from pre- to post-sleep.

We also measured memory changes from pre- to post-sleep cued recall tasks. Two independent raters rated identification, detail, and gist from the cued recall task according to previous studies on the verbal recall of emotional scenes (Catarino et al., 2015), if there was inconsistent between the two raters, another rater would be involved to reconcile the discrepancies. Memory change scores were calculated by subtracting pre-sleep baseline memory scores from post-sleep memory scores, with higher change scores indicating larger memory retention.

### ERPs and time-frequency analyses

For ERPs, artifact-free short epochs were averaged, and baseline corrected (pairing trial: -1s to 0s; non-pairing trial: -1 s to 0s). For time-frequency analysis, a continuous wavelet transformation with variance cycles (3 cycles in length at 1 Hz, increasing linearly with frequency to 15 cycles at 30 Hz) was implemented on pairing trial epochs (-1.5s to 5.5 s) and non-pairing trial epochs (-1.5s to 3.5s) to obtain power for the frequency range from 1 to 30 Hz, in steps of 0.5 Hz and 5ms. Epochs were cropped to eliminate edge artifacts (pairing trial: -1s to 5s; non-pairing trial: -1s to 3s) after time-frequency transformation. Subsequently, averaged spectral power was normalized (Z-scored) using a [-1 to -0.2 s] baseline for the pairing trial and for the non-pairing trial, separately.

We reported time-frequency and ERP results from the pairing trials to investigate the neural mechanisms of affective updating. For non-pairing trials that only involved cues, we hypothesized that EEG activity may capture the online change of positive vs. neutral vs. non-pairing memory cues. However, we did not find differences between these conditions. Results of non-pairing block were reported in the Supplementary (S4).

### Sleep staging analysis

We conducted sleep stage scoring based on a machine learning algorithm Yet Another Spindle Algorithm (YASA, Vallat & Walker, 2021), was double-checked by an experienced sleep researcher. EEG data were first re-referenced to FPz per YASA recommendations. The C4 (or C3 if C4 was marked as a bad channel), EOG, and EMG channels were used to feed the algorithm. Before statistics on sleep staging could be calculated, artifacts had to be identified. Table S1 provides information on sleep stages.

### Slow oscillations and spindle detection

We extracted slow oscillations (SOs) and sleep spindles implemented in YASA (Vallat & Walker, 2021). SOs were detected at Fz based on previous research (e.g., Helfrich et al., 2017; Mölle et al., 2002). EEGs were first bandpass filtered (0.5-2 Hz) using a FIR filter with a transition band of 0.2 Hz. Second, after zero-crossings were detected, events were selected based on duration (0.5s-2s) and amplitude (75 percentile) criteria. Individual SOs were detected on each trial from the [-15 to 15s] 30s long epochs, with the detection results retained in the [-1.5 to 5.5s] 7s epochs.

Sleep spindles were detected at Cz (Schechtman et al., 2021), using the root mean square (RMS). EEGs were first down-sampled to 100 Hz, followed by bandpass filtered between 11 and 16 Hz. Second, the RMS was calculated at every sample point with a sliding window of 300 ms at a step of 100 ms. Spindles thresholds were determined by the mean of RMS plus 1.5 SDs of the signals. The 10% lowest and 10% highest values were removed before computing the SD of RMS. If a sample exceeds this threshold, it would be tagged as a potential spindle. Next, for neighboring potential spindles, they were merged together if the between-spindles intervals were shorter than 500 ms. Spindle events were counted only if they met the 0.5s-2s duration criterion. Spindles were detected on each [-15 to 15s] 30s long epoch, with the detection results retained in the [-1.5 to 5.5s] 7s epochs.

### SO phase analysis

To investigate how temporal coupling between the word onset and SO phase influences affective updating, we conducted an item-level analysis focusing on the SO phase when playing positive pairing cues. We divided the positive pairing cues into negative-change vs. negative-stay sub-conditions based on pre- to post-sleep affect judgment changes. We defined trials as negative-change when the affect judgments changed from pre-sleep negative to post-sleep neutral, i.e., successful affective updating. We defined trials as negative-stay when both pre- and post-sleep affect judgments were negative, i.e., no affective updating. Participants were excluded from this analysis if they did not have negative-change trials. In the positive pairing condition, 26 subjects were retained. To examine whether the effect was specifically due to positive pairing, we repeated this analysis with neutral pairing cues, with 25 subjects retained.

We next examined the SO phase clustering of negative-change and negative-stay trials in positive and neutral pairing conditions, separately. The SOs were identified using the method described above. Given that the word-cue pairing occurred during NREM sleep, we used trials with at least two SOs between the -1.5s and 5.5s for subsequent phase analysis. To eliminate the biases of different trial numbers between negative-change and negative-stay sub-conditions, we matched the trial number by retaining the temporally closest trials in these two sub-conditions. Next, we extracted the instantaneous phase of the onset of spoken words and memory cues using a Hilbert transform. We examined the coupling between word/cue onset and SO phases using the Rayleigh test and V test. Specifically, the Rayleigh test examines non-uniformity of event distributions, with a significant result indicating that the events are preferably clustered toward certain phase angles and thus followed a non-uniform distribution. The V test examines whether the clustering would occur at a pre-specified phase angle (e.g., 0°; peak), against uniform distributions or the clustering would occur at a different phase angle than the pre-specified phase.

To further validate the robustness of the SO phase effect, we conducted an inverted analysis. First, upon detection of SOs in each pairing trial, we assigned the trial to two sub-conditions: emotional words upstate vs. downstate, pairing cues upstate vs. downstate, according to whether their onsets were located between the mid crossing of a SO and its end (upstate) or between the start of a SO and its mid crossing (downstate). We counted the number of trials in each sub-condition, and conducted a linear mixed model to explore whether the number of trials in these conditions influenced affective updating. We first focused on the upstate number of the emotional words, using the formula described below:

Affective updating ∼ 1 + emotional_words_upstate *condition + (1 + emotional_words_upstate | subject).

‘emotional_words_upstate’ was a continuous variable, denoting the number of emotional words delivered during the SO upstate. ‘condition’ was a categorical variable (positive vs. neutral paring).

Next, we focus on the SO upstate trial number of the memory cue. The formula was as follows:

Affective updating ∼ 1 + memory_cue_upstate*condition + (1 +memory_cue_upstate | subject).

‘memory_cue_upstate’ was a continuous variable, denoting the number of memory cues delivered at the SO upstate.

## Results

### Sleep pairing updated affective judgment but not memory recall

To answer our primary research question on sleep-based affective updating, we examined affect-judgment changes from pre- to post-sleep. In the affect-judgment task, we calculated the neutral response ratio by dividing the number of neutral responses by the number of trials in each of the three conditions. At the pre-sleep learning session, we confirmed that emotional learning was successful, such that participants were more likely to judge the cues as negative than neutral: *t*(30) = -14.43, *p* < .001, ***d*** = 2.59. Moreover, there was no significant difference between conditions in neutral response ratio during the pre-sleep learning session (Mean ± S.E.; positive pairing: 0.41 ± 0.048; neutral pairing: 0.46 ± 0.048; non-pairing: 0.38 ± 0.045; *F*(2,60) = 2.21, *p* = .118).

To quantify the affect-updating effect, pre-sleep neutral response ratio was subtracting from the post-sleep neutral response ratio to calculate the affect-change score, we used this affect-change score to measure affective updating. We found that there was a significant difference between positive pairing, neutral pairing, and non-pairing conditions (*F*(2,60) = 4.23, *p* = .030, **η**^**2**^ = 0.12). Post-hoc tests revealed a higher affect-change score for the positive pairing compared to neutral pairing (Mean ± S.E., positive pairing: 0.07± 0.02; neutral pairing: 0.01± 0.02; *t*(30) = 2.36, *p* = .037, FDR corrected, ***d*** = 0.46) and to non-pairing (non-pairing: 0.01± 0.03; *t*(30) = 2.41, *p* = .037, FDR corrected, ***d*** = 0.42). We did not observe a significant difference between neutral pairing and non-pairing (*t*(29) = 0.10, *p* = .92).

We next tested whether RTs in the affect-judgment task differed by condition. A 3 (positive pairing vs. neutral pairing vs. non-pairing) * 2 (negative vs. neutral response) repeated-measures ANOVA was conducted on RT changes from pre- to post-sleep. There were no significant differences for condition, valence, nor their interaction (*p*s>.19). The same analyses on subjective valence and arousal rating changes did not reveal significant main nor interaction effects (*p*s>.62).

We next sought to explore whether our procedure produced changes in the recall of negative memories. Identification and gist changes were calculated by dividing pre-sleep correct responses by post-sleep correct responses. Memory detail scores were Z-normalized within participants to control the variance of participants’ verbal descriptions (Zhuang et al., 2021). Then, memory detail change scores were calculated by subtracting pre-sleep from post-sleep memory detail scores. There were no significant differences among the three conditions on these three memory changes scores (Gist: *F*(2,60) = 0.75, *p* = .479, **η**^**2**^ = 0.02, Identification: *F*(2,60) = 0.17, *p* = .840, **η**^**2**^ = 0.01; Detail: *F*(2,60) = 0.29, *p* = .752, **η**^**2**^ = 0.01).

### Spoken words during NREM sleep elicited ERPs 450 ms following word onset

To demonstrate that the sleeping brain responded to spoken words, we first calculated auditory evoked brain potentials across all electrodes. The butterfly plot revealed EEG responses peaked around 450 ms after word onset (see Figure 2A). A time-series of whole-brain responses to spoken words were computed using global field power (Figure 2B). Given that we played two stimuli (word+cue) in a pairing trial, we found two peaks after the onset of each stimulus, one at 450 ms and another at 2450 ms). We analyzed corresponding ERP amplitudes across all electrodes, averaging artifact-free epochs across all trials following 1-s pre-stimulus baseline-correction. A permutation *t*-test was performed across electrodes to compare ERPs to zero; Figure 2A illustrates significant electrodes (*p*s < .049). These results suggest that the sleeping brain responded to both auditory word stimuli.

**Figure 1:**
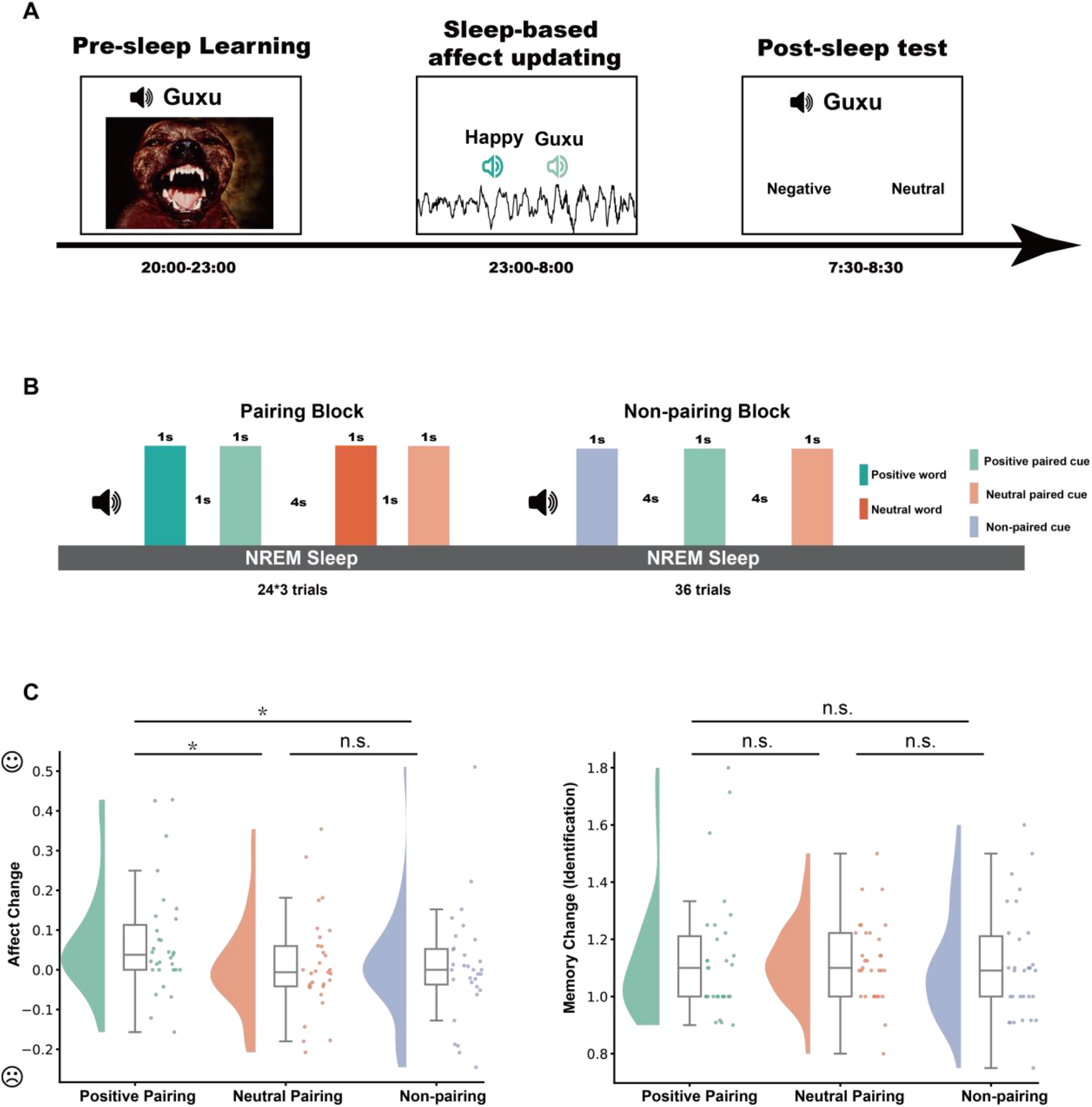
Experiment procedure and overnight affective updating. (A) During pre-sleep learning, participants memorized 36 pseudowords + negative picture pairs. During subsequent NREM sleep, the experimenter played spoken words and cues to participants (until ∼ 2 am or until 7 rounds were completed, whichever came earlier). After waking in the morning, participants completed post-sleep tests including the affect-judgment task and the cued verbal-recall task. An example trial of the affect-judgment task is shown. (B) Sleep-based affective-updating procedure during sleep: three sleep pairing blocks and one non-pairing block constitute one round. Each pairing block consisted of 24 trials (12 positive word+cue pairings and 12 neutral word+cue pairings). The non-pairing block included 36 trials, including the same 24 cues from the preceding pairing blocks as well as the remaining 12 cues from pre-sleep learning. (C) Behavioral outcomes of affective updating (left) and memory changes (right). *: *p* < .05, **: *p* < .01.

**Figure 2:**
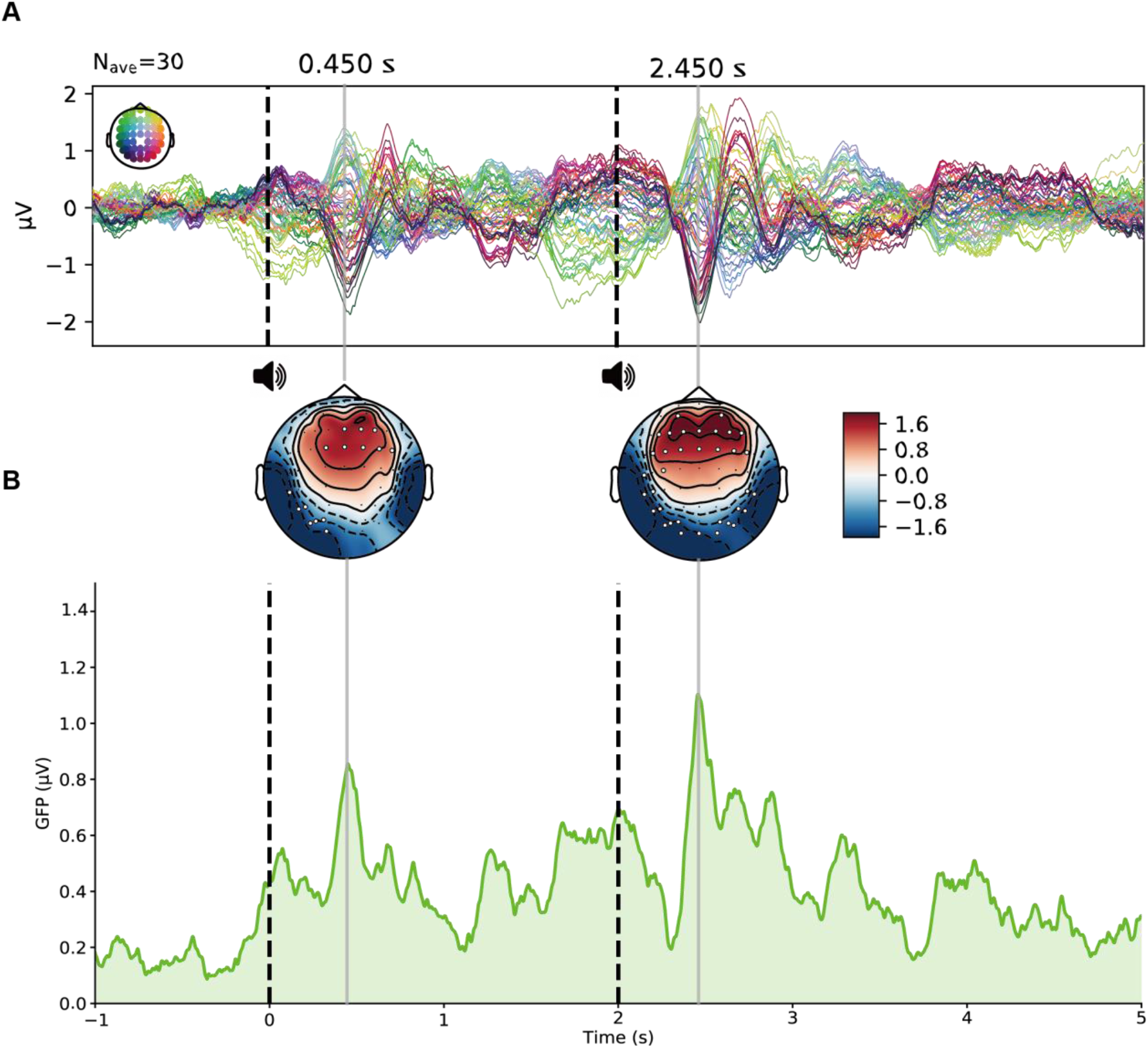
ERPs elicited by spoken words during NREM sleep. (A) Butterfly plot of ERP to the spoken words collapsing across positive and neutral pairing conditions. (B) The Global field power (GFP) plot revealed two peaks at 450 ms after word onset. At each time point, GFP was the standard deviation of all electrodes. The topographical plot displayed the significant electrodes of ERP at two peaks when comparing the ERP to zero.

We were also interested in whether ERPs differed between the positive pairing and neutral pairing conditions. A permutation *t*-test was also performed to assess differences between positive and neutral pairings at the two peaks. The results revealed no statistically significant differences in ERPs between positive and neutral pairings (*p*s>.455).

### Spoken words elicited the delta-theta and sigma-beta power during NREM sleep

To investigate stimulus-elicited EEG activity, we performed time-frequency analysis on EEG epochs followed by averaging across conditions and participants (Figure 3A). Via a nonparametric permutation test across time points and frequency bands at Cz, we identified three positive clusters and one negative cluster, which showed that sound playing significantly modulates the earlier delta-theta-alpha cluster (1-12Hz) and later sigma-beta cluster (11-25Hz) (Clusters *p*s < 0.019; Figure 2D). We first focused on EEG responses elicited by the emotional words (positive vs. neutral) within two clusters (delta-theta-alpha: 0.36s-1.07s; and sigma-beta: 0.59s-1.84s). The memory cue also elicited two clusters (delta-theta-alpha: 2.30s-3.04s; sigma-beta: 2.37s-3.84s). We focused our analysis on the positive clusters. These positive clusters were generally consistent with previous TMR or sleep-learning studies that reported that stimuli (auditory tones or spoken words) modulated brain activity during sleep (Schechtman et al., 2021; Züst et al., 2019). We then used these identified clusters as regions of interest (ROIs) in the following analysis.

**Figure 3:**
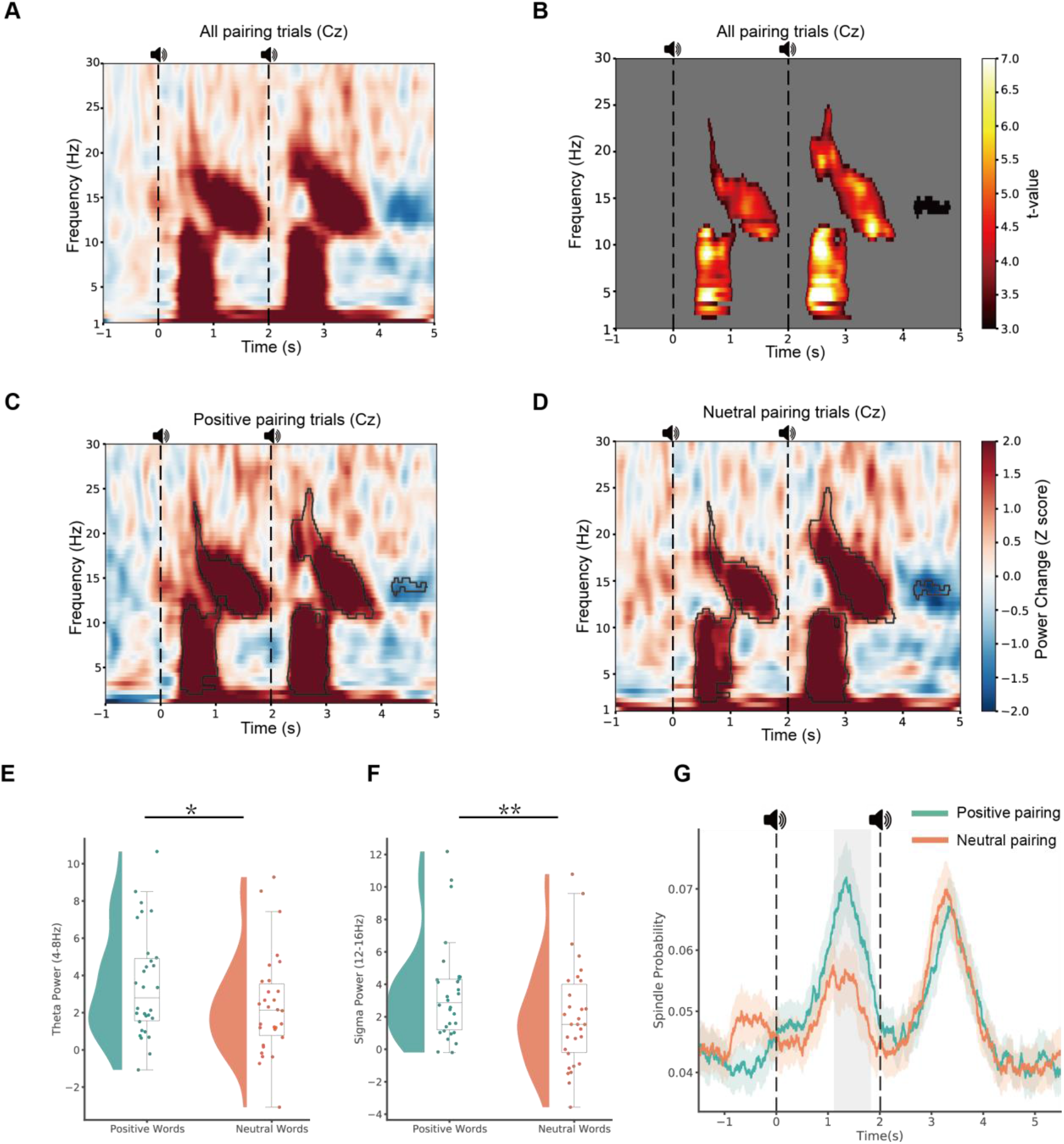
Stimulus-elicited EEG activity and spindle probability. (A) Time-frequency results of auditory processing during sleep averaged over all trials and subjects at Cz. (B) A cluster-based permutation test across frequency bands and time points at Cz results in a t-values map for auditory stimulus modulating neural oscillations during NREM sleep. A time-frequency plot for positive (C) and neutral (D) pairing conditions, blackline highlighting significant cluster area. Difference in theta (E) and sigma (F) power (from the significant cluster between positive and neutral words). (G) Spindle probability difference between positive and neutral pairings over time, shaded area indicates SE. *: *p* < 0.05, **: *p* < 0.01.

### Emotional valence modulated theta and spindle/sigma activity during NREM sleep

To examine whether the sleeping brain would distinguish between positive and neutral spoken words, we directly compared the EEG power elicited by positive and neutral words within the significant clusters identified in the abovementioned analyses. The results showed that that positive words elicited a significantly larger power increase than neutral words across delta, theta, and alpha band (Mean ± S.E., Positive word: 3.31 ± 0.28; Neutral word: 2.22 ± 0.27; *t*(29)=2.30, *p* = .030, 95% CI[0.12, 2.06], ***d*** = 0.46). To further delineate the frequency-specific effect, we focused on delta (1-4Hz) and theta (5-9Hz), according to previous studies (Canales-Johnson et al., 2019; Legendre et al., 2022; Lehmann et al., 2016). The results showed that positive words elicited significantly stronger theta power than neutral words (Figure 3E, theta: *t*(29) = 2.25, *p* = .033, 95% CI[0.11, 2.27], ***d*** = 0.44), while no significant effect was observed in the delta band (*t*(29) = 1.625, *p* = .115).

We also examined the effect of emotional valence sigma-beta range (12-25Hz) activity as identified in the above clusters during NREM sleep. A paired t-test showed that positive words elicited a significantly larger power increase than neutral words (Mean ± S.E., Positive word: 3.30 ± 0.31; Neutral word: 1.96 ± 0.32; *t*(29) = 2.79, *p* = .009, 95% CI[0.36, 2.31], ***d*** = 0.44), More specifically, we found that positive words elicited significantly greater sigma power than neutral words (*t*(29) = 2.82, *p* = .009, 95% CI: [0.38, 2.40], ***d*** = 0.439). However, this effect was not observed in the beta band (*t*(29) = 1.552, *p* = .131). These results suggest that word valence modulates theta and sigma power change during NREM sleep.

To further understand whether observed sigma effects were driven by discrete spindle activity, we examined spindle number in the different conditions. An automatic spindle-detection technique (see Methods) was used on single trials to determine the spindle probability at each time point of the trial (Schechtman et al., 2021). We tested whether positive words induced a higher spindle probability than neutral words. A permutation test was conducted on spindle probability across time. We found that positive words elicited a higher spindle probability than neutral words from 1130-1810 ms post-stimulus (Figure 3G, *p*_cluster_ = .021).

### Theta and spindle/sigma activity differed in response to the paired stimuli

We next asked whether pairing valence and pairing position modulated theta and sigma power. We conducted a 2 (pairing valence: positive vs. neutral pairing) * 2 (pairing position: emotional words vs. memory cues) repeated-measures ANOVA on theta and sigma power separately within the corresponding significant clusters.

Regarding theta power, we found a significant main effect of pairing position (Figure 4A, *F*(1,29) = 9.77, *p* = .004, **η**^**2**^ = 0.25), indicating that the memory cue elicited a larger theta power change than the emotional word (Mean ± S.E., memory cue: 3.55 ± 0.49 vs. emotional word: 2.74 ± 0.42). However, both the main effect of pairing valence (*F*(1,29) = 3.17, *p* = .086) and the interaction (*F*(1,29) = 1.38, *p* = .249) were not significant. This effect was replicated when using the whole delta-theta cluster but was not when using the delta power, suggesting the effect is driven by theta activity (see supplementary S1).

**Figure 4:**
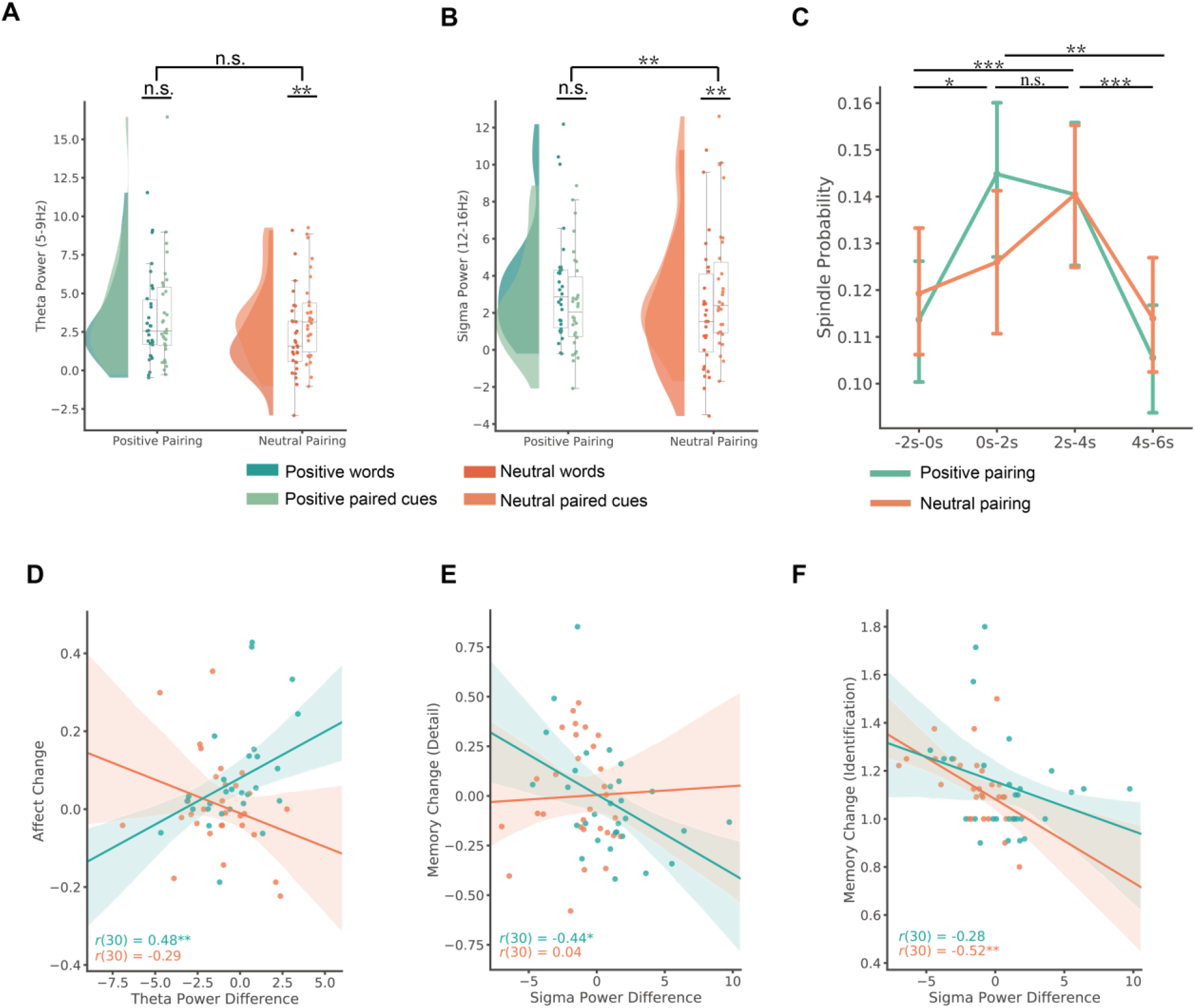
The effect of pairing valence and position on theta, sigma, and spindle probabilities. (A). Theta power: memory cues induced a significantly larger theta power than emotional words, irrespective of the valence. (B). Sigma power: positive paired cues elicited a similar sigma power increase to the positive words, whereas neutral pairing cues elicited a significantly larger sigma power increase than neutral words. (C). Spindle probability at every 2-second interval during the pairing trial, with error bar indicating 95% CI. (D). Theta power differences in positive words and positive paired cues positively predicted affective updating. (E). Sigma power differences in positive words and positive paired cues negatively predicted the detail of memory change. (F). Sigma power differences in neutral words and neutral paired cues negatively predicted the identification of memory change. *: *p* < .05, **: *p* < .01, ***: *p* < .001, shaded area indicates 95% CI.

Regarding sigma power, the same analyses did not find a significant main effect of pairing valence (Figure 4B, *F*(1,29) = 0.53, *p* = .471) nor a main effect of pairing position (*F*(1,29) = 0.35, *p* = .560). However, the valence by position interaction was significant (*F*(1,29) = 10.13, *p* = .003, **η**^**2**^ = 0.26). Post-hoc comparisons showed that in the neutral pairing condition, the neutral word elicited a lower sigma power increase than the paired cue (Mean ± S.E., neutral word: 2.05 ± 0.60; cue: 3.35 ± 0.63; *t*(29) = -3.27, *p* = .006, FDR corrected, 95% CI[-2.11, -0.49], ***d*** = 0.39); whereas in the positive pairing condition, the positive word elicited slightly higher sigma power increase than the paired cue (Mean ± S.E., positive word: 3.43 ± 0.56; paired cue: 2.52 ± 0.49; *t*(29) = 1.67, *p* = .106). The results were consistent when using the entire sigma-beta band, while there were no significant effects when analyses focused on beta band (see supplementary S2).

We next examined how pairing valence modulated spindle probability following the two stimuli along time. We summed spindle probability for every 2 s and conducted a repeated-measures ANOVA with valence (Positive pairing, Neutral pairing) and time intervals (-2s-0, 0-2s, 2s-4s,4s-6s), note there would be 1s overlap between the 4-6s of the current trial and the -2-0s of the following trial, given that each epoch is -1.5s – 5.5s long. Although the main effect of pairing was not significant (*F*(1,29) = 0.31, *p* = .583), the main effect of time intervals (*F*(3,87) = 12.87, *p* < .001, **η**^**2**^ = 0.307) and the interaction were significant (*F*(3,87) = 3.27, *p* = .034, see Figure 4C). Decomposing the pairing valence by time interval interaction, we found that positive words elicited significantly higher spindle probabilities than neutral words during the 0-2s interval (*t*(29) = 2.70, *p* = .046, FDR corrected); while no significant differences were found during the other intervals (*t*s(29) < 0.71, *p*s > .767).

The significant time interval effect was driven by enhanced spindle activity shortly after playing the emotional word (0-2s, Mean ± S.E., 0.14 ± 0.007) and after the memory cues (2-4s, 0.14 ± 0.008), when compared to pre-stimulus baseline (0.12 ± 0.01) and the 4-6 s late interval (0.11 ± 0.006). Detailed statistics of pairwise comparisons are provided in supplementary S6.

### Theta power difference between positive words and memory cues predicted affective updating

We next sought to ask whether the theta and sigma power change implicated in sleep pairing has any effect on affective updating. To quantify the sleep pairing effect at a neural level, we subtracted power induced by memory cues from the power induced by emotional words. This subtraction removed the non-specific auditory processing, and captured EEG power differences between emotional words and memory cues. A higher value would indicate stronger neural processing of emotional words than memory cues, and possibly more effective affective conditioning effect.

Using this metric, we next examined the relationship between theta and sigma power difference with the affective updating, respectively. The significant correlation was observed in the theta power difference (Figure 4D, *r*(30) = 0.48, 95%CI [0.15, 0.72], *p* = .007) but not in the sigma power difference (*r*(30) = 0.12, *p* = .528) in the positive pairing condition. However, no significant correlations observed in the neutral pairing conditions (theta: *r*(30) = -0.29, *p* = .126; sigma: *r*(30) = 0.14, *p* = .466). The correlation between EEG power difference and affective updating became 0.71 when using the whole cluster (see supplementary S3).

To verify that the prediction effect was driven by the theta power differences between the positive words and the positive paired cues, and not by the power elicited by either single word, we re-ran the analyses using partial correlation to control the power elicited by single positive words and positive paired cues. Results remained significant after controlling for theta power elicited by each single word in the pairing (theta: *r*(30) = 0.47, CI: [0.12, 0.72], *p* = .011). Therefore, the theta power difference between the positive words and the following cues significantly predicted overnight affective updating.

### Sigma power difference between emotional words and memory cues predicted memory change

Spindle-related sigma power has been linked to memory processing during sleep. In TMR, while post-cue sigma power positively predicted memory consolidation, pre-cue sigma showed opposite predictions (Antony, Cheng, et al., 2018; Antony, Piloto, et al., 2018; Wang et al., 2019). We thus asked whether the sigma power differences between the emotional words and the paired cues can predict memory changes before and after sleep. We found that all three memory change scores (identification, gist, details) were negatively correlated with the sigma power differences. though we only found a significant negative correlation between the sigma power difference and the memory change of detail in the positive pairing conditions (Figure 4E, *r*(30) = -0.44, 95%CI [-0.69, -0.10], *p* = .015) and the memory change of identification in the neutral pairing conditions (Figure 4F, *r*(30) = -0.52, CI: [-0.74, -0.19], *p* = .003). Partial correlation confirmed that only the sigma power difference, rather than the sigma power induced by either single word, predicted memory change (detail in positive pairing condition: *r*(30) = -0.46, 95%CI [-0.71, -0.10], *p* = .015; identification in the neutral pairing condition: *r*(30) = -0.50, 95%CI [-0.74, -0.16], *p* = .007). Together, these results suggested that during pairing, stronger sigma power elicited by the emotional words relative to memory cues would result in more forgetting of negative memories following sleep.

### Successful affective updating depends on positive-word onset within an SO upstate

Recent sleep learning and TMR studies suggest that the precise coupling between SO upstates and cueing contributes to successful sleep encoding and TMR (Batterink et al., 2016; Göldi et al., 2019; Züst et al., 2019). We are thus also interested in examining the relationship between SO-event coupling and affective updating. To quantify successful affective updating at an item level, we sub-grouped trials into negative-change and negative-stay trials based on performance in the affect judgment task (see Methods). We next collapsed negative-change trials across all participants and extracted the SO phase when emotional words and cues were played. In the positive pairing condition, we found that negative-change trials were associated with a significant non-uniform distribution of positive word onsets (*Z*(681) = 7.46, *p* < .001, Rayleigh test) and of the following cues (*Z*(681) = 4.87, *p* = .008, Rayleigh test). In addition, we found that the onset of positive words (V test against 0°: *v* = 71.02, *p* < .001, mvl = 0.10; coupling phase: -4.91°, circular mean) and the onset of positive paired cues (V test against 0°: *v* = 57.41, *p* < .001, mvl = 0.08; coupling phase: -4.17°, circular mean) were both preferentially coupled to the SO peak (i.e., upstate). However, in the negative-stay condition, the onset of positive words and positive pairing cues were randomly distributed (Positive words: *Z*(681) = 1.75, *p* = .174; Positive paired cues: *Z*(681) = 1.34, *p* = .263, Rayleigh tests). We next conducted the same analysis in the neutral pairing condition, and did not find significant clustering in the negative-change trials (Neutral words: *Z*(412) = 0.57, *p* = .567; Neutral paired cues: *Z*(412) = 0.61, *p* = .541, Rayleigh test) or in the negative-stay trials (Neutral words: *Z*(412) = 1.20, *p* = .30; Neutral paired cues: *Z*(412) = 0.37, *p* = .690, Rayleigh test). Our SO phase results indicated that for affective updating to be successful, that is for participants to judge memories as more neutral due to positive pairing, the onset of positive words and cues were both coupled to the SO peak. Note that the phase effect was specific to SO events, as the same analyses using the delta band (2-4Hz) did not yield significant effects (see supplementary S4). Moreover, when we conducted the SO phase analysis at the participant level, results did not change (see supplementary S5).

Next, we conducted an inverted analysis to confirm the robustness of our phase result. We used the linear mixed model to explore whether the number of trials (of either emotional words or memory cues onset) delivered during an SO upstate modulated affective updating. Regarding the emotional words, we found a significant main effect of pairing condition (χ^2^(1) = 9.13, *p* = .003) and interaction effect between pairing condition and upstate trial number (χ^2^(1) = 4.25, *p* = .039). Post-hoc comparison revealed a stronger association between upstate trial number and positive pairing than the association in the neutral pairing condition (*b* = 0.025, SE = 0.012, *t*(688) = 2.05, *p* = .040), indicating the more positive words delivered at the SO upstate, the larger the affective change following sleep (Figure 5C, left panel).

**Figure 5:**
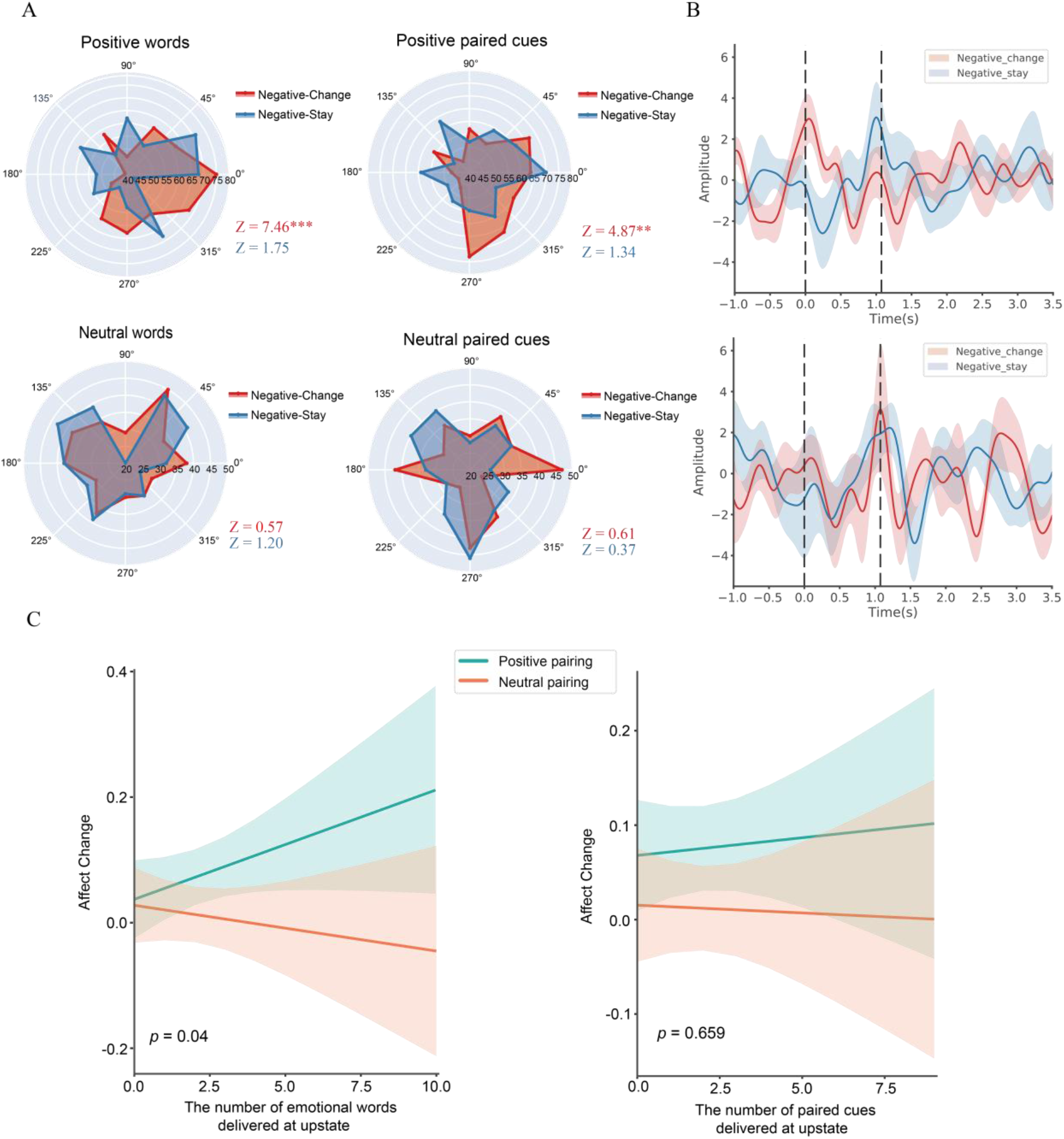
Relationship between slow oscillation phase and word onset. (A) The onset phase of slow oscillation for positive words and paired cues from successful affective updating (negative-change) or not (negative-stay). Negative-change and negative-stay trials were combined across all subjects, and the Rayleigh test was used to examine the phase distribution of each condition. Negative-change trials were significantly non-uniformly distributed during the onset of positive words and paired cues. (B) Grand average ERPs from negative-change and negative-stay in positive pairings (upper panel) and neutral pairings (lower panel), with a low-pass filter at 2 Hz applied. (C) The number of positive words delivered at the SO upstate modulated successful affective updating, with shaded area indicating 95% CI. **: *p* < .01 ***: *p* < .001

For the memory cue, we found that the main effect of pairing condition was significant (χ^2^(1) = 8.77, *p* = .003). However, we did not find any significant effect on upstate trial number (χ^2^(1) = 0.022, *p* = .883) and their interaction (χ^2^(1) = 0.197, *p* = .657, Figure 5C right panel). Taken together, these SO phase analyses indicated that when positive words were coupled with SO upstate, affective updating was more successful.

## Discussion

Can unwanted memories be updated during sleep, when people can avoid the impact of recalling a negative life event? We demonstrated that via pairing positive words with memory cues during NREM sleep, participants’ affect judgments became less negative, which we term affective updating. In addition to this behavioral effect, we found that greater theta power increases to positive words than to memory cues predicted successful affective updating. In contrast, greater sigma power to the positive word than to the cue predicted forgetting. Notably, at an item-level, the timing of positive word onset to a slow oscillation upstate contributed to successful affective updating. By demonstrating a sleep-based affective updating effect with associated neural correlates, the present study provides important knowledge to guide new possibilities for editing unwanted memories.

Despite the apparent disconnection from the external world, the sleeping brain responds to external stimuli with a preserved information-processing capacity, as evidenced by stimulus-elicited theta and spindle activity. Specifically, emotional prosody, tone, memory cue, and even relaxing words could modulate theta power during sleep (Beck et al., 2021; Blume et al., 2017; Canales-Johnson et al., 2019; Lehmann et al., 2016). In addition, auditory processing can modulate spindle-related sigma power (Andrillon et al., 2016; Andrillon & Kouider, 2020; Wislowska et al., 2022). Consistent with this research, we showed that the emotional valence conveyed by words modulated theta and spindle/sigma activity, which was associated with memory updating as discussed below.

Observing that emotional words modulate theta and sigma activity, how exactly might this neural activity be involved in affective updating? Given that theta power induced by positive words could indicate affective information processing, and theta power induced by memory cues may track reactivation of emotional memories (Legendre et al., 2022; Lehmann et al., 2016; Schreiner et al., 2017), we postulated that theta differences between emotional words and memory cues could reflect something about the memory modulation. Accordingly, we quantified the pairing effect by calculating the theta power differences between emotional words and memory cues. Our results indicated that larger the theta power elicited by positive words than memory cues, the more affective updating was shown. However, there was no such relationship in the neutral pairing condition. Thus, successful affective updating may depend on theta activity elicited by positive words, implicating affective encoding during sleep.

In terms of memory changes, while we did not find a main effect of valence pairing, it is worth noting that sigma power difference between the emotional word and memory cue predicted forgetting. Intriguingly, we found significant interactions between pairing valence (positive vs. neutral trials) and pairing position (emotional word vs. memory cue) on both sigma power and spindle probabilities. Specifically, the emotional words elicited stronger sigma power and higher spindle probabilities than the neutral words, while such differences became weaker for the memory cues. Moreover, the temporal trajectory of spindle probability (Figures 3G and 4C) was consistent with spindle refractory hypothesis, such that spindles are segregated by refractory periods, and a second spindle would be less likely to occur within 3-6s after the first spindle (Antony, Piloto, et al., 2018). Regarding sigma power and memory reprocessing, previous studies showed that pre-cue sigma negatively predicted post-cue sigma power as well as the TMR-induced memory consolidation (Antony, Cheng, et al., 2018; Antony, Piloto, et al., 2018; Wang et al., 2019). In our study, given that the two stimuli were played consecutively, sigma induced by the emotional words could function as pre-cue sigma preceding the subsequent cues, with larger pre-cue sigma power suppressing post-cue sigma power. Accordingly, stronger sigma power to the emotional word relative to the memory cue modulated memory consolidation and induce forgetting.

Temporal coupling between the external stimuli and SO upstates has been shown to be conducive for successful sleep encoding and memory reactivation (Göldi et al., 2019; Züst et al., 2019). Indeed, SO upstates represent unique periods associated with cortical excitability and neural plasticity that may be essential for information processing during sleep (Destexhe et al., 2007; Schabus et al., 2012). Corroborating this hypothesis of the SO upstate, our results found that at an item-level, successful affective updating depended on coincidence between words onset and SO upstate. Scrutinizing the coupling results suggested that the onset of positive words, but not memory cues, drove affective updating. These results complement a recent sleep learning study, which showed that successful sleep learning occurred when the second word of the word pairings was delivered at the SO peaks (Züst et al., 2019). Unlike sleep learning wherein a novel word was paired with a known word and participants learnt novel semantic associations, our paradigm involved pairing of positive words and memory cues, or counterconditioning (Hu et al., 2017; Keller et al., 2020). Extending sleep learning research, our study showed that optimal processing of the positive stimuli, as indicated by higher theta power and precise coupling with SO upstate, was crucial to update the affect of the associated memory.

Limitations and future directions shall be noted. First, while we included non-pairing blocks to examine whether we could capture the online neural representation change of the memory cues due to pairing, we did not find such evidence. Notably, Arzi et al., (2012) found that nasal airflow and delta-theta activity could capture the online sleep affect learning effect. This discrepancy might be due to the emotional word used in our study being less potent than the pleasant/aversive odor used in Arzi et al., (2012). Future studies might examine the effectiveness of different sensory modalities (e.g., auditory vs. olfactory) in memory updating during sleep. Second, whether the sleep-based affective updating effect can be long-lasting remains unknown, given that we did not include a delayed test. Future studies may examine the long-term effect of sleep pairing in updating unwanted memories. Third, while the affective updating effect is evident in the affect judgment task that captured spontaneous and fast affect responses, subjective emotional ratings did not show such updating effects. Previous research suggests that sleep learning and TMR effects are more evident using indirect measures such as nasal airflow, response speed, and forced choice tasks (Arzi et al., 2012, 2014; Cairney et al., 2014; Hu et al., 2015; Koroma et al., 2022; Züst et al., 2019). Future research should clarify the extent to which the sleep learning benefits are evident in different behavioral tasks.

During sleep, the brain continues processing sensory stimuli despite ostensible disconnection from the external world (Andrillon & Kouider, 2020). Harnessing the power of the sleeping brain, we showed that responses to memory cues could be changed via pairing positive words with these cues during NREM sleep. We further identified cardinal sleep EEG signals such as theta and sigma activity, as well as the coupling between emotional stimuli and SO upstates, that played instrumental roles supporting emotion and memory dynamics. The present study provides insights into how to develop novel paradigms to update or modify unwelcomed memories, and pinpoints possible neural mechanisms supporting effective updating. An important question that remains to be tackled in future research will be how to help people better manage unwanted memories they have acquired outside the laboratory, such as from actual traumatic experiences.

## Acknowledgements

This research was supported by the National Natural Science Foundation of China (No. 31922089, No.321791056) and General Research Fund (No. 17601318) of Hong Kong Research Grants Council to X.H., and Grants NSF BCS-2048681and NIH/NINDS R01 NS112942 to K.A.P.

## Author contributions

Tao Xia: Conceptualization, Methodology, Data curation, Writing- Original draft preparation. Software, Project administration. Ziqing Yao.: Writing - Review & Editing, Visualization. Xue Guo: Data curation, Writing - Review & Editing. Jing Liu: Writing - Review & Editing. Danni Chen: Writing - Review & Editing. Qiang Liu: Data curation, Resources. Ken A. Paller: Writing - Review & Editing, Funding acquisition. Xiaoqing Hu: Conceptualization, Methodology, Writing- Original draft preparation. Writing- Reviewing and Editing, Funding acquisition, Supervision.

## Data and code Availability

Preprocessed data and the code used for analysis are available online via Github after publication.

## Conflict of interest statement

None

## Supplementary analyses

### S1 Delta-theta-alpha and sigma-beta cluster differed in response to the emotional words and the memory cues

Besides theta power, we also conducted a 2(positive pairing vs. neutral pairing) *2(first vs. second) repeated ANOVA analyses on the delta power band and the whole cluster. Regarding the delta band power, we did not find a significant effect on the main effect of sleep pairing (*F*(1,29) = 1.28, *p* = .267) and pairing position (*F*(1,29) = 3.79, *p* = .061) and their interaction (*F*(1,29) = 2.12, *p* = .156).

Regarding the whole cluster, we found that there was a significant main effect of pairing position (*F*(1,29) = 13.05, *p* = .001, **η**^**2**^ = 0.31), post-hoc tests showed that the memory cue elicited a greater delta-theta-alpha power change than the emotional words (Mean ± S.E., emotional words: 2.76 ± 0.26; memory cues: 3.47 ± 0.26 ; *t*(29) = -3.61, *p* = .001, 95% CI: [-1.11, -0.31], ***d*** = 0.36). However, we did not observe a significant effect on the main effect of sleep pairing (*F*(1,29) = 2.30, *p* = .14, **η**^**2**^= 0.07), and on the interaction (*F*(1,29) = 3.56, *p* = .07, **η**^**2**^ = 0.11). The results from whole cluster across delta-theta-alpha analysis were consistent with results of theta analysis in the main results.

Besides sigma power, we also conducted a 2(positive pairing vs. neutral pairing) *2(first vs. second) repeated ANOVA analyses on the beta power band and the whole cluster. Regarding the beta band power, we did not find a significant effect on the main effect of sleep pairing (*F*(1,29) = 0.71, *p* = .407) and pairing position (*F*(1,29) = 0.70, *p* = .409) and their interaction (*F*(1,29) = 4.14, *p* = .051).

Regarding the sigma-beta cluster, neither the main effect of sleep learning (*F*(1,29) = 1.08, *p* = .31) nor the pairing position (*F*(1,29) = 0.14, *p* = .71) were not significant. However, the interaction effect of sleep learning and pairing position was significant (*F*(1,29) = 11.44, *p* = .002, **η**^**2**^ = 0.28). Post-hoc comparisons showed that positive words elicited greater sigma-beta power change than paired cues (Mean ± S.E., Positive words: 3.30 ± 0.53; Paired cues: 2.20 ± 0.38; *t*(29) = 2.30, *p* = .029, FDR corrected, 95% CI[0.36, 2.31], ***d*** = 0.43), whereas the neutral words elicited smaller sigma power change than the paired cues (Mean ± S.E., Neutral words: 1.96 ± 0.57; Paired cues: 2.84 ± 0.49; *t*(29) = 2.63, *p* = .027, FDR corrected, 95% CI[0.36, 2.31], ***d*** = 0.30). The results from whole cluster across sigma-beta analysis were consistent with results of sigma analysis in the main results, and further indicating that the relationship between emotional words and memory cues had reverse pattern in positive and neutral pairs.

### S2 Delta-theta-alpha and sigma-beta cluster power difference between emotional words and memory cues predicted affective updating

When using the whole cluster, a significant positive correlation was also observed between the delta-theta-alpha cluster power change and the affect change score in positive pairing (*r*(30) = 0.71, CI: [0.48, 0.85], *p* < .001) but not in neutral pairing conditions (*r*(30) = -0.16, CI: [-0.50, 0.21], *p* = .385), In addition, no significant correlation delta power change and affect change was found in the positive (*r*(30) = 0.35, *p* = .059) and neutral pairings (*r*(30) = 0.11, *p* = .581).

We also correlated the sigma-beta cluster power difference with the memory change before and after sleep. In the positive learning condition, we found that sigma-beta power difference could significantly predict the memory change of detail (*r*(30) = -0.40, CI: [-0.66, -0.04], *p* = .030) but cannot predict the memory change of gist (*r*(30) = -0.18, CI: [-0.50, 0.2], *p* = .354) and identification (*r*(30) = -0.20, CI: [-0.52, 0.17], *p* = .289). In the neutral learning condition, however, the sigma-beta power difference can predict memory change of identification (*r*(30) = -0.46, CI: [-0.71, 0.10], *p* = .010) but cannot predict the memory change of detail (*r*(30) = -0.007, CI: [-0.37, 0.35], *p* = .971) and gist (*r*(30) = -0.13, CI: [-0.47, 0.23], *p* = .503). In addition, we did not observe a significantly correlation between the beta power difference and memory change of detail in positive pairings (*r*(30) = -0.12, *p* = .525) and did not observe a significantly correlation between the beta power difference and memory change of identification in neutral pairings (*r*(30) = -0.27, *p* = .143). These results suggested that the effects of power difference predicted affect and memory change were specific to theta and sigma band.

### S3 Control analysis for slow oscillation phase analysis

Regarding the SO phase analysis in negative-change and negative-stay for positive-pairing trials, we did a control analysis by using the phase of the delta band (2-4Hz). In the negative-change trials from all participants and the preferred phase of each participant, we found the delta phase distribution at the onset of positive words and positive pairing cues were randomly distributed (*Z*s < 2.30 *p*s > .100).

### S4 ERP results of non-pairing block

We conducted the same analysis of pairing blocks on non-pairing blocks. The butterfly plot revealed responses to words around 450ms after the word onset. A time series of whole-brain responses to memory cues were computed using global field power. We found one peak after playing memory cues (450 ms). We analyzed corresponding ERPs amplitudes across all electrodes, averaging the artifact-free epochs across all trials following 1 s pre-stimulus baseline correction,. The permutation t-test was performed across electrodes at the peak to compare the ERPs with zero. Results showed that there were no significant channels higher than zero.

We were also interested in whether there was a difference in ERP between the positive pairing and neutral pairings. A permutation t-test was also performed to assess the differences between positive and neutral pairings at the two peaks. The results revealed no statistically significant difference in ERP between the positive and neutral pairings (*p*s> .455).

#### Time-frequency analysis

The logic of time-frequency analysis in the non-pairing block was the same as in the pairing block. We first run a permutation test across time points and frequency bands at Cz. Three positive clusters were identified across the delta, theta, alpha, and sigma bands (*p*_clusters_< .005). These clusters were then considered as regions of interest in the next analysis. Power values within each band in the identified cluster were extracted from positive paired, neutral paired, and non-paired cues. We did not find any difference among the three conditions across interested power bands (*F*_s_(2,56) = 0.94, *p*_s_ > .395).

### S5 SO phase analyses at a participant level

To test the robustness of these SO phase results, we conducted similar analyses at a participant level, complementing item-level analyses reported in the main texts. In the positive pairing condition, we still observed that for each participant, the averaged preferred phase of negative-change trials were coupled to the SO peak (Positive words: *Z*(26) = 6.18, *p* = .002, Rayleigh test; V test against 0°: *v* = 12.02, *p* < .001, mvl = 0.49; coupling phase: - 18.51°, circular mean; Positive paired cues: *Z*(26) = 2.98, *p* = .049, Rayleigh test; V test against 0°: *v* = 8.74, *p* = 008, mvl = 0.34; coupling phase: -6.54°, circular mean).

**Table S1.** Time spent in each sleep stage.

|  | TIB (min) | N1 (min) | N2 (min) | N3 (min) | NREM (min) | REM (min) |
| --- | --- | --- | --- | --- | --- | --- |
| Mean | 504 | 28 | 203 | 79 | 310 | 63 |
| SEM | 3.3 | 3.2 | 7.9 | 7.1 | 11.5 | 5.7 |

### S6 Pairwise t-tests for spindle probability between time intervals

Inspecting Figure 4C suggested that spindle probability during post-stimulus 0-2s and 2-4s were significantly higher than the pre-stimulus baseline -2s-0s and post-stimulus 4-6s. This observation was confirmed statistically (Mean ± S.E. p-values were FDR corrected): 0-2s vs. -2-0s: 0.14 ± 0.007 vs. 0.12 ± 0.01, *t*(29) = 2.89, *p*_corrected_ = .010, ***d*** = 0.52; 0-2s vs. 4-6s: : 0.14 ± 0.01 vs. 0.11 ± 0.006, *t*(29) = 3.80, *p*_corrected_ = .001, ***d*** = 0.71; 2-4s vs. -2-0s: 0.14 ± 0.008 vs. 0.12 ± 0.006, *t*(29) = 4.75, *p*_corrected_ < 0.001, ***d*** = 0.64; 2-4s vs. 4-6s: 0.14 ± 0.008 vs. 0.11 ± 0.006, *t*(29) = 4.67, *p*_corrected_ < .001, ***d*** = 0.82.

## Notes

### Competing Interest Statement

The authors have declared no competing interest.

